# Sequential sequestrations increase the incorporation and retention of multiple growth factors in mineralized collagen scaffolds

**DOI:** 10.1101/2020.05.11.089524

**Authors:** Aleczandria S. Tiffany, Marley J. Dewey, Brendan A.C. Harley

## Abstract

Trauma induced injuries of the mouth, jaw, face, and related structures present unique clinical challenges due to their large size and complex geometry. Growth factor signaling coordinates the behavior of multiple cell types following an injury, and effective coordination of growth factor availability within a biomaterial can be critical for accelerating bone healing. Mineralized collagen scaffolds are a class of degradable biomaterial whose biophysical and compositional parameters can be adjusted to facilitate cell invasion and tissue remodeling. Here we describe the use of modified simulated body fluid treatments to enable sequential sequestration of bone morphogenic protein 2 and vascular endothelial growth factor into mineralized collagen scaffolds for bone repair. We report the capability of these scaffolds to sequester growth factors from solution without additional crosslinking treatments and show high levels of retention for individual and multiple growth factors that can be layered into the material via sequential sequestration steps. Sequentially sequestering growth factors allows prolonged release of growth factors *in vitro* and suggests the potential to improve healing of large-scale bone injury models *in vivo*. Future work will utilize this sequestration method to induce cellular activities critical to bone healing such as vessel formation and cell migration.

## 1. Introduction

Craniomaxillofacial injuries – injuries of the mouth, jaw, face, and related structures – can be caused by a wide range of congenital abnormalities, oral cancer treatments, and traumatic injuries. Trauma related injuries experienced by civilians^1–5^ and high-energy impact injuries experienced by Warfighters^6–8^ present unique clinical challenges. These injuries are often large, complex in geometry, and cannot be repaired with external fixtures alone^9,10^. Most clinical treatments use autografts, bone taken from a secondary site in the patient with the injury, or allografts, bone taken from a human donor^11–13^. While autografts are considered the gold-standard and maintain osteo-conductive and osteo-inductive abilities, allografts are a popular alternative because of their availability^14,15^. However, there are limitations to the use of these bone grafts. Autografts are limited by the size of the injury site, and allografts raise concerns about disease transmission, transplant rejection, and their purification methods are not uniform, resulting in variability between grafts^16–18^. Thus, there is a clinical need for alternative solutions to address critically sized craniomaxillofacial injuries.

Regenerative medicine solutions commonly seek to combine a biomaterial template with strategies to accelerate healing such as the incorporation of biomolecule stimuli^19,20^. Bone healing is a multistep process with multiple cell types and is coordinated through growth factor signaling^21–25^. Briefly, inflammatory cytokines such as tumor necrosis factor alpha are released following injury and lead to the formation of a hematoma^26,27^. These cytokines recruit macrophages and other immune cells to the injured site and initiate new vessel formation^27^. Vascular endothelial growth factor (VEGF) and angiopoietin-1 and 2 are critical growth factors for promoting and maintaining angiogenesis^26,28–30^. Immune cells then release factors such as bone morphogenic protein 7 to recruit mesenchymal stem cells to the injury site^31^. Mesenchymal stems cells start depositing extracellular matrix components such as collagen that lead to callous formation at the injured site^22,24^. Bone morphogenic protein 2 (BMP2) and other signaling molecules induce the differentiation of mesenchymal stem cells into osteoblasts (bone depositing cells) and mineralization of the callous occurs^31,32^. As the injury continues to heal, growth factors such as osteoprotegerin recruit osteoblasts and osteoclasts to form mechanically weak bone^33^. This weak bone will continue to be remodeled and will eventually be fully replaced by mechanically robust bone^25^.

Due to the complexity of native bone healing, a wide range of efforts have explored the use of these and other factors to accelerate cell recruitment and regenerative activity. For example, stromal derived factor 1 and platelet derived growth factor have been delivered from bone-mimetic scaffolds to increase cell migration *in vitro*^34^ and improve bone healing *in vivo*^35,36^. Incorporating BMP2 into scaffolds for bone repair is very common and has shown to improve osteogenesis in multiple systems^36–42^. However, the need for doses of soluble BMP2 larger than what’s normally found in the body has led to complications such as abnormal bone formation^43,44^. Prolonged release of VEGF from bone-mimetic scaffolds in small concentrations has been shown to promote vascularization of the injury site and improve bone healing *in vivo*^45^. However, delivery strategies that result in quick release of high concentrations of VEGF increase vascularization at the expense of bone quantity *in vivo*^46^. Thus, it is critical to determine the optimal dose for growth factor delivery, understanding that this dose will vary based on the desired results, and develop strategies to control the delivery and release of factors from biomaterial substrates to accelerate healing.

Our lab has developed a mineralized collagen-glycosaminoglycan scaffold capable of inducing osteogenesis without the addition of exogenous factors^47^, and these materials have been used to heal sub-critical injuries *in vivo*^48,49^. However, as we move into larger injury models, we may require biomolecular supplements to improve implant-bone integration, cell recruitment, and vascular remodeling. Thus, we are interested in exploring how growth factor supplementation can be used to improve *in vivo* healing and integration with host tissue. Our group has previously used strategies such as photopatterning^50,51^, covalent immobilization^52–55^, and sequestration modulated by glycosaminoglycan content^56^ and small molecules^57^ to add growth factors to non-mineralized and mineralized collagen scaffolds. Recently, the Murphy lab has described the use of modified simulated body fluid (mSBF) to create mineral coatings on material surfaces to deliver growth factors^58–62^, plasmid DNA lipoplexes^63^, and condensed mRNA^64^. We were particularly interested in the work by Clements *et al*. in which they performed sequential sequestrations to layer Interleukin-1 Receptor Antagonist on nanoparticles to prolong *in vivo* activity^62^.

In this manuscript, we describe the use of mSBF and sequential sequestration to incorporate and retain BMP2 and VEGF in three-dimensional mineralized collagen scaffolds. First, we hypothesized that soaking mineralized collagen scaffolds in mSBF prior to sequestration would increase incorporation and retention of growth factors within our scaffolds. Next, we hypothesized that sequential sequestrations would increase incorporation and retention of growth factors within our scaffolds compared to a single sequestration. We examined the capacity for mineralized collagen scaffolds to sequester growth factors from solution without additional crosslinking treatments and the capability to increase incorporation and extend retention of single growth factors or multiple growth factors via sequential sequestrations.

## 2. Materials and Methods

### 2.1. Mineralized collagen scaffold fabrication

Mineralized collagen-glycosaminoglycan scaffolds were fabricated via lyophilization from a mineralized collagen precursor suspension as described before^47,65,66^. The precursor suspension was created by homogenizing type I collagen (1.9 weight per volume, Sigma Aldrich, St. Louis, Missouri USA), chondroitin-6-sulfate (0.84 weight per volume, Sigma Aldrich), and calcium salts (calcium hydroxide and calcium nitrate, Sigma Aldrich) in a mineral buffer solution (0.1456M phosphoric acid/0.037M calcium hydroxide). The precursor suspension was stored at 4°C and degassed prior to lyophilization.

Mineralized collagen scaffolds were fabricated via lyophilization using a Genesis freeze-dryer (VirTis, Gardener, New York USA) as described before^66^. Briefly, 100uL of precursor suspension was pipetted into a custom 144-well polysulfone mold (6mm diameter, 7mm tall wells). The precursor solution was frozen by cooling from 20°C to -10°C at a constant rate of 1°C/minute followed by a temperature hold at -10°C for 2 hours. The frozen suspension was then sublimated at 0°C and 0.2 Torr, resulting in a porous scaffold network.

All scaffolds were hydrated for 2 hours in ethanol, crosslinked for 2 hours in EDC-NHS, and washed in phosphate buffered saline (PBS) for 48 hours prior to use in experiments.

### 2.2. Modified simulated body fluid preparation

Modified simulated body fluid (mSBF) was made according to previous recipes^58–64^. Briefly,1.41 mM sodium chloride (NaCl, Sigma Aldrich), 4.0 mM potassium chloride (KCl, Sigma Aldrich), 0.5 mM magnesium sulfate (MgSO_4_, Sigma Aldrich), 1.0 mM magnesium chloride (MgCl_2_, Sigma Aldrich), 5.0 mM calcium chloride (CaCl_2_, Sigma Aldrich), 1.0 mM potassium phosphate (KH_2_PO_4_, Sigma Aldrich), and 4.2 mM sodium carbonate (NaHCO_3_, Sigma Aldrich) were dissolved in deionized water and sterile filtered through a MilliporeSigma Stericup with a 0.22 μm filter (Fisher Scientific, Hampton, New Hampshire USA). mSBF was stored at 4°C until use.

### 2.3. Sequestration of bone morphogenic protein 2 following modified simulated body fluid treatments

Following hydration, scaffolds were soaked in 50 ng/mL bone morphogenic protein 2 (BMP2) or treated in modified simulated body fluid (mSBF) for 1, 3, or 7 days and then soaked in 50 ng/mL BMP2 (**Figure 1A**). mSBF soaks were done at room temperature under mild shaking and mSBF solution was changed each day. BMP2 was diluted in 1% bovine serum albumin in phosphate buffered saline (1% BSA in PBS), and sequestration was done for 1 hour at room temperature under mild shaking. After sequestration, the BMP2 solution was saved and stored at -20°C and scaffolds were placed in PBS for 7 days at 37°C. PBS was replaced each day and stored at - 20°C. An enzyme-linked immunosorbent assay (ELISA) (R&D Systems, Minneapolis, Minnesota, USA) was used to quantify the amount of BMP2 sequestered and retained within the scaffolds. Retention is reported as the percent of BMP2 remaining in the scaffold to BMP2 initially sequestered into the scaffold.

**Figure 1:**
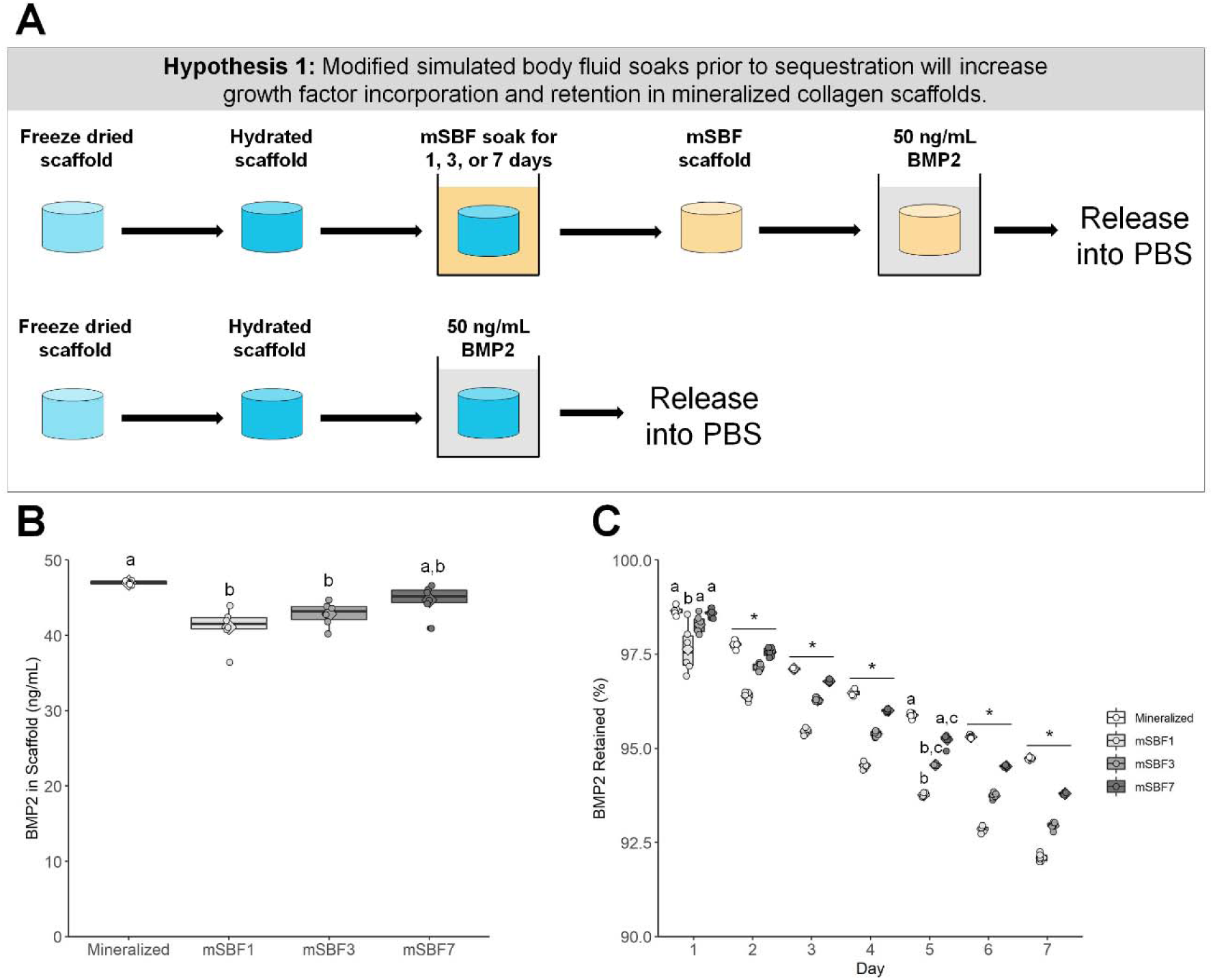
Growth factor sequestration with varying modified simulated body fluid (mSBF) treatments. (A) Schematic of experiment. Hydrated scaffolds and scaffolds soaked in mSBF for 1, 3, or 7 days were used to sequester BMP2. (B) Sequestration of BMP2 after varying treatment times in mSBF. Mineralized collagen scaffolds are capable of sequestering BMP2 without mSBF treatments. Groups that share a letter are not significantly different (p<0.05). (C) Retention of BMP2 within scaffolds for 7 days. Mineralized collagen scaffolds had the highest retention compared to all mSBF treated groups by Day 7. Groups that share a letter within a day are not significantly different (p<0.05). * indicates significant differences between all groups within a day. *NOTE: The y-axis starts at 90% to better show differences between groups. To see plots with the y-axis 0-100% please see Supplemental Figure 1.*

### 2.4. Compression testing

Stress-strain curves of hydrated scaffolds, scaffolds soaked in phosphate-buffered saline (PBS) for 7 days, and scaffolds soaked in modified simulated body fluid (mSBF) for 7 days were generated with the Instron 5943 mechanical tester (Instron, Norwood, Massachusetts USA) using a 5 N load cell; hydrated scaffolds were not submerged in liquid during testing. Samples were compressed at a rate of 1 mm/min with the Young’s Modulus determined from the stress-strain curves using conventional analysis methods for low-density open-cell foam structures such as the mineralized collagen scaffolds^67,68^.

### 2.5. Sequential sequestration of single growth factors

#### 2.5.1. One treatment

Following hydration, scaffolds were soaked in 80 ng/mL bone morphogenic protein 2 (BMP2) or vascular endothelial growth factor (VEGF) (**Figure 1A**). Growth factors were diluted in 1% bovine serum albumin in phosphate buffered saline (1% BSA in PBS), and sequestration was done for 1 hour at room temperature under mild shaking. After sequestration, the growth factor solution was saved and stored at -20°C and scaffolds were placed in PBS for 7 days at 37°C.

#### 2.5.2. Sequential treatments

Following hydration, scaffolds were soaked in 10 ng/mL BMP2 or VEGF and put into modified simulated body fluid (mSBF) overnight at room temperature under mild shaking. This was repeated for 7 consecutive days (8 total sequestrations) (**Figure 2A**). Growth factors were diluted in 1% BSA in PBS, and sequestration was done for 1 hour at room temperature under mild shaking. After each sequestration, the growth factor solution was saved and stored at - 20°C. The solution from mSBF soaks were saved and stored at -20°C. After the final sequestration, scaffolds were placed in PBS for 7 days at 37°C.

**Figure 2.**
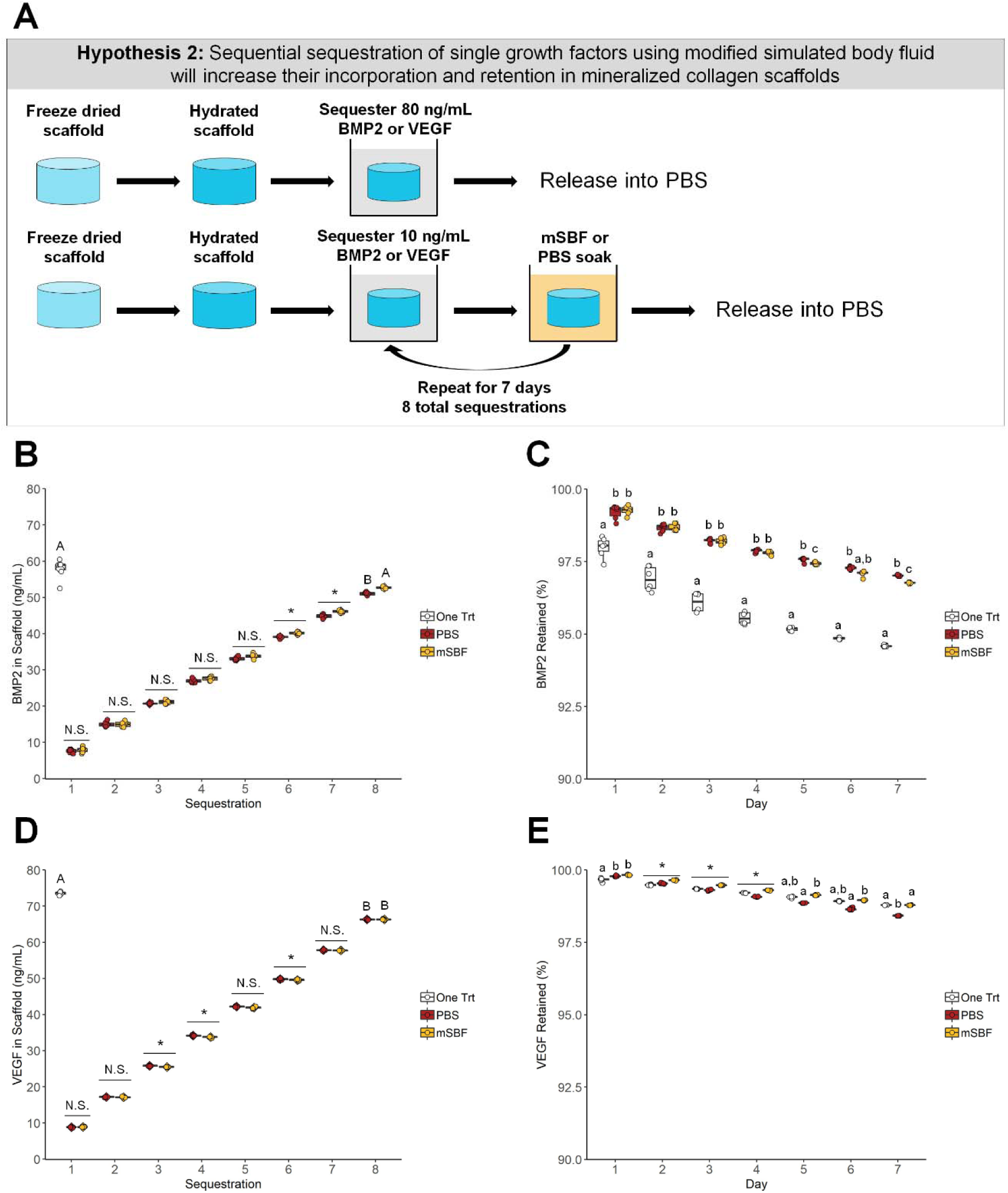
Sequential sequestration of individual growth factors. (A) Schematic of experiment. Scaffolds soaked once in BMP2 or VEGF were compared to sequentially sequestered BMP2 or VEGF groups. (B) BMP2 sequestration. The one-time BMP2 treatment and the modified simulated body fluid (mSBF) sequentially sequestered groups had the same amount of BMP2 sequestered (p<0.05). * indicates significance between all groups within a day (p<0.05). N.S. indicates no significance between groups within a day (p<0.05). Groups that share a letter are not significantly different (p<0.05). (C) BMP2 retention. Scaffolds treated once with BMP2 had lower retention of the growth factor in the scaffold compared to both sequentially sequestered groups. Groups that share a letter within a day are not significantly different (p<0.05). *NOTE: The y-axis starts at 90% to better show differences between groups. To see plots with the y-axis 0-100% please see Supplemental Figure 3.* (D) VEGF sequestration. The one-time VEGF treatment had more VEGF sequestered than both sequentially sequestered groups (p<0.05). * indicates significance between groups within a day (p<0.05). N.S. indicates no significance between groups within a day (p<0.05). Groups that share a letter are not significantly different (p<0.05). (E) VEGF retention. Scaffolds soaked once in VEGF and scaffolds sequentially sequestered with mSBF treatments have the highest retention by Day 7. * indicates significance between all groups within a day (p<0.05). Groups that share a letter within a day are not significantly different (p<0.05). *NOTE: The y-axis starts at 90% to better show differences between groups. To see plots with the y-axis 0-100% please see Supplemental Figure 3*.

### 2.6. Sequential sequestration of multiple growth factors

#### 2.6.1. One treatment

Following hydration, scaffolds were soaked in 80 ng/mL bone morphogenic protein 2 (BMP2) and 10 ng/mL vascular endothelial growth factor (VEGF) (**Figure 3A**). Growth factors were diluted in 1% bovine serum albumin in phosphate buffered saline (1% BSA in PBS), and sequestration was done for 1 hour at room temperature under mild shaking. After sequestration, the growth factor solution was saved and stored at -20°C and scaffolds were placed in PBS for 7 days at 37°C.

**Figure 3.**
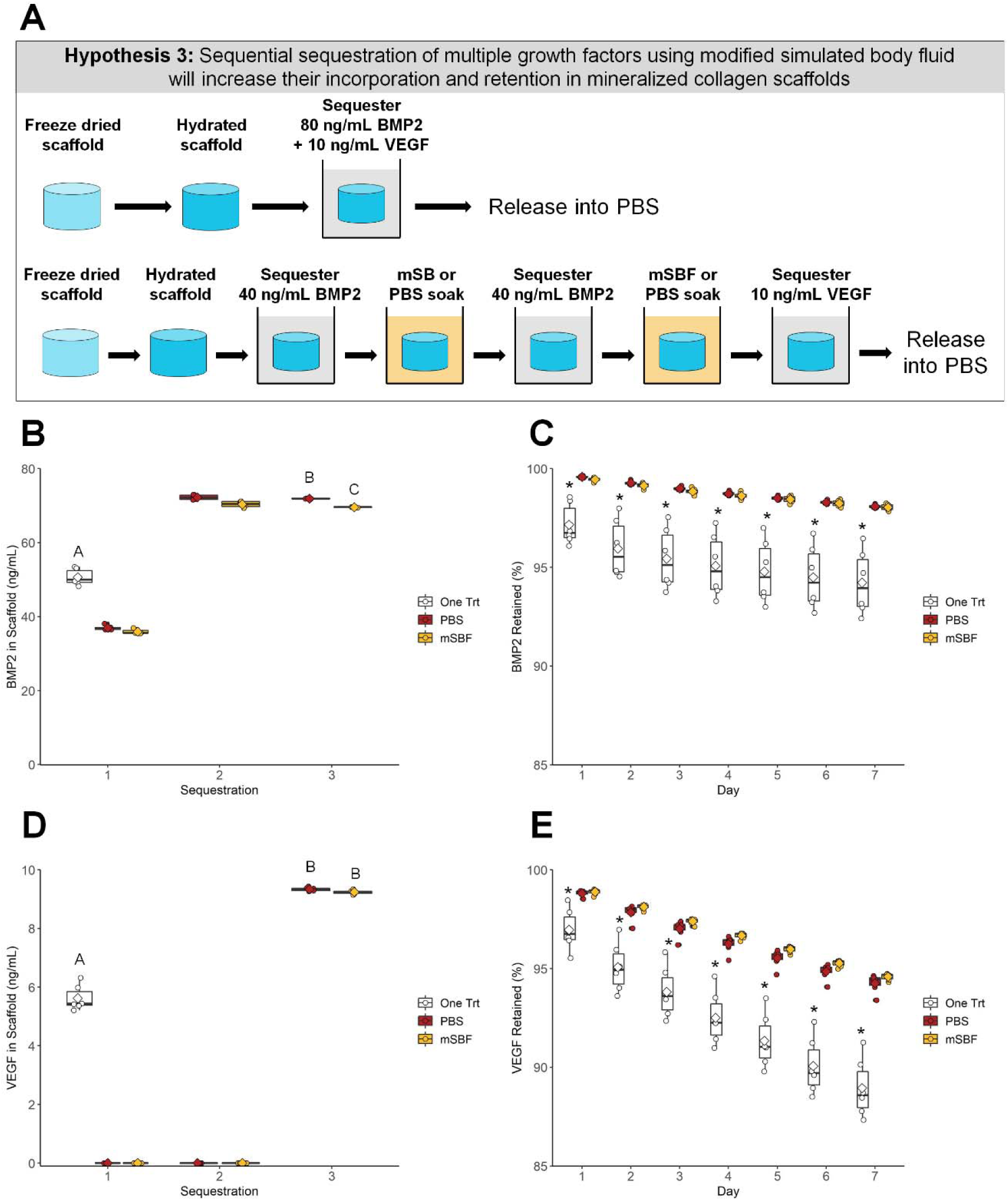
Sequential sequestration of multiple growth factors. (A) Schematic of experiment. Scaffolds soaked once in a solution of BMP2 and VEGF were compared to sequentially sequestered BMP2 then VEGF groups. (B) BMP2 sequestration. Both sequentially sequestered scaffolds had higher concentrations of BMP2 compared to the one-treatment scaffolds. Groups that share a letter are not significantly different (p<0.05). (C) BMP2 retention. Scaffolds treated once in BMP2 have the lowest retention compared to both sequentially sequestered groups (p<0.05). * indicates significant differences compared to the other two groups within a day (p<0.05). *NOTE: The y-axis starts at 85% to better show differences between groups. To see plots with the y-axis 0-100% please see Supplemental Figure 4.* (D) VEGF sequestration. Both sequentially sequestered scaffolds had higher concentrations of VEGF compared to the one-treatment scaffolds. Groups that share a letter are not significantly different (p<0.05). (E) VEGF retention. Scaffolds treated once in VEGF have the lowest retention compared to both sequentially sequestered scaffolds. * indicates significant differences compared to the other two groups within a day (p<0.05). *NOTE: The y-axis starts at 85% to better show differences between groups. To see plots with the y-axis 0-100% please see Supplemental Figure 4.*

#### 2.6.2. Sequential treatments

Following hydration, scaffolds were soaked twice in 40 ng/mL BMP2 and once in 10 ng/mL VEGF with overnight modified simulated body fluid (mSBF) soaks following each sequestration step (**Figure 2A**). mBSF soaks were done at room temperature under mild shaking. Growth factors were diluted in 1% BSA in PBS, and sequestration was done for 1 hour at room temperature under mild shaking. After each sequestration, the growth factor solution was saved and stored at -20°C. The solution from mSBF soaks were saved and stored at -20°C. After the final sequestration, scaffolds were placed in PBS for 7 days at 37°C.

### 2.7. Enzyme-linked immunosorbent assays

#### 2.7.1. One-time treatments

PBS was replaced each day and stored at -20°C during growth factor release. An enzyme-linked immunosorbent assay (ELISA) (R&D Systems) was used to quantify the amount of BMP2 and VEGF sequestered and retained within the scaffolds. Retention is reported as the percent of BMP2 remaining in the scaffold to BMP2 initially sequestered into the scaffold.

#### 2.7.2. One-time treatments

PBS was replaced each day and stored at -20°C during growth factor release. An enzyme-linked immunosorbent assay (ELISA) (R&D Systems) was used to quantify the amount of BMP2 and VEGF sequestered and retained within the scaffolds. Growth factor released during mBSF soaks was factored into the final concentration of BMP2 or VEGF sequestered into mineralized collagen scaffolds. Retention is reported as the percent of BMP2/VEGF remaining in the scaffold to BMP2/VEGF initially sequestered into the scaffold.

### 2.8. Human umbilical vein endothelial cell culture and preliminary transwell experiment

Human umbilical vein endothelial cells (HUVECs) (Lonza, Basel, Switzerland) were expanded in T75 flasks (Fisher Scientific) and cultured in endothelial cell growth media (Lonza) at 37°C and 5% carbon dioxide until confluent. Once confluent, passage 4 HUVECs were seeded (62,000 cells/well) into the lower chamber of the transwell plate and scaffolds were loaded into the transwell insert. 300uL of endothelial cell growth media was used in the lower chamber and 700uL of endothelial cell growth media was used in the transwell insert. Cells were cultured in endothelial cell growth media at 37°C and 5% carbon dioxide for 7 days.

Four groups were used for this preliminary transwell experiment: (1) scaffolds soaked in PBS with no VEGF added into the transwell insert (Blank); (2) scaffolds soaked in PBS with 10 ng/mL VEGF added into the transwell insert (Soluble); (3) scaffolds that had been soaked in 5 ng/mL of VEGF, soaked in mSBF overnight, and soaked in another 5 ng/mL of VEGF (mSBF); (4) scaffolds soaked once in 10 ng/mL VEGF (One Trt). The Soluble groups had no additional VEGF added after the first media change at Day 3.

### 2.9. Human umbilical vein endothelial cell metabolic activity

The metabolic activity of human umbilical vein endothelial cells (HUVECs) seeded in transwells was measured using alamarBlue via fluorescent spectrophotometer (Tecan Infinite F200 Pro, Männedorf, Switzerland). HUVECs seeded in the bottom chamber of a transwell were incubated in a 10% alamarBlue solution (Invitrogen, Carlsbad, California USA) for 90 minutes at 37°C under moderate shaking. The relative cell metabolic activity was determined from a standard curve generated with known cell concentrations. An experimental value of 1 indicates the metabolic activity of the number of cells originally seeded into the transwell.

### 2.10. Human umbilical vein endothelial cell gene expression

RNA was isolated from cell seeded scaffolds using TRIzol Reagent (Invitrogen) following the provided protocol. Isolated RNA was quantified using the NanoDrop Lite Spectrophotometer (ThermoFisher, Waltham, Massachusetts USA) and reverse transcribed using a QuantiTect Reverse Transcription Kit (Qiagen, Hilden, Germany) and a BioRad S1000 thermal cycler (BioRad, Hercules, California USA).

Each real-time polymerase chain reaction (PCR) was carried out in duplicate, using 10 ng cDNA and Taqman primers (ThermoFisher). A Taqman PCR kit (ThermoFisher) along with an Applied Biosystems 7900HT Fast Real-Time PCR System was used to perform the real-time PCR. 18S ribosomal RNA (18S) and peptidylprolyl isomerase A (PPIA) were used as housekeeping genes. Gene expression profiles were obtained for platelet derived growth factor (PDGF), hypoxia inducible factor 1 alpha (Hif1a), angiopoietin 1 (Ang1), and angiopoietin 2 (Ang2) (details: **Supplemental Table 1**).

### 2.11. Statistics

RStudio was used for all plotting (ggplot2) and statistical analysis. No outliers were removed. Sample size was six (n=6) for all experiments except compression testing (n=12) and polymerase chain reaction (PCR, n=8). PCR data have some groups with sample size less than 8 due to undermined threshold cycle values, but 90% of groups have n≥6. See **Supplemental Tables 2 and 3** for details on sample size for PCR data. Data presented in all tables are presented as average ± standard deviation.

#### 2.11.1. Data with two experimental groups

A t-test was used for data that were normally distributed and had equal variance. A Welch’s t-test was used for data that were normally distributed and did not have equal variance. The Mann-Whitney U Test was used for data that were not normally distributed and had equal variance. Experimental groups were tested for normality using the Shapiro-Wilk test. Homogeneity of variance was tested using the Levene Test.

#### 2.11.2. Data with three or more experimental groups

The Kruskal-Wallis test (one experimental factor) was used for data that were not normally distributed and had equal variance, and significance was determined using Dunn’s post-hoc test. One-way ANOVA (one experimental factor) or two-way ANOVA (two experimental factors) was run for data that were normally distributed and had equal variance, and significance was determined using Tukey’s post-hoc test. Welch’s one-way ANOVA was used for data that were normally distributed and did not have equal variance, and significance was determined using Tukey’s post-hoc test. Mood’s Median Test (via a Monte Carlo simulation, one experimental factor) was used for data that were not normally distributed and did not have equal variance, and significance was determined using a pairwise median test. Residuals were tested for normality using the Shapiro-Wilk test. Homogeneity of variance was tested using the Levene Test.

## 3. Results

### 3.1. Mineralized collagen scaffolds can sequester growth factors without additional treatments

We first determined whether extended soaking of mineralized collagen scaffolds in modified simulated fluid (mSBF) prior to growth factor sequestration would increase incorporation and retention of growth factors within our scaffolds. Our mineralized collagen scaffold natively sequesters bone morphogenic protein 2 (BMP2) better than mineralized collagen scaffolds soaked in mSBF for 1 or 3 days (**Figure 1B**). However, extended exposure to mSBF (7 days) recovers this sequestration ability, with no significant difference in BMP2 sequestration compared to the native mineralized scaffolds (**Figure 1B**). All groups sequestered greater than 80% of BMP2 out of solution (**Figure 1B**). The mineralized collagen scaffolds also retained the highest percentage of sequestered BMP2 over 7 days compared to all mSBF soaked scaffold groups, with the difference becoming significant by Day 2 (**Figure 1C**). Overall, mineralized collagen scaffolds had the highest final concentration of BMP2 at Day 7 than scaffolds with extended mSBF treatments prior to sequestration (**Table 1**). Scaffolds that were soaked in mSBF for 7 days both sequestered and retained BMP2 better than shorter mSBF exposures (1 and 3 days); however, subsequent experiments did not include mSBF pre-soaks because there was no improvement in BMP2 sequestration or retention compared to native mineralized collagen scaffolds. Summary statistics and full-scale retention data for this experiment: **Table 1, Supplemental Figure 1**.

**Table 1:**
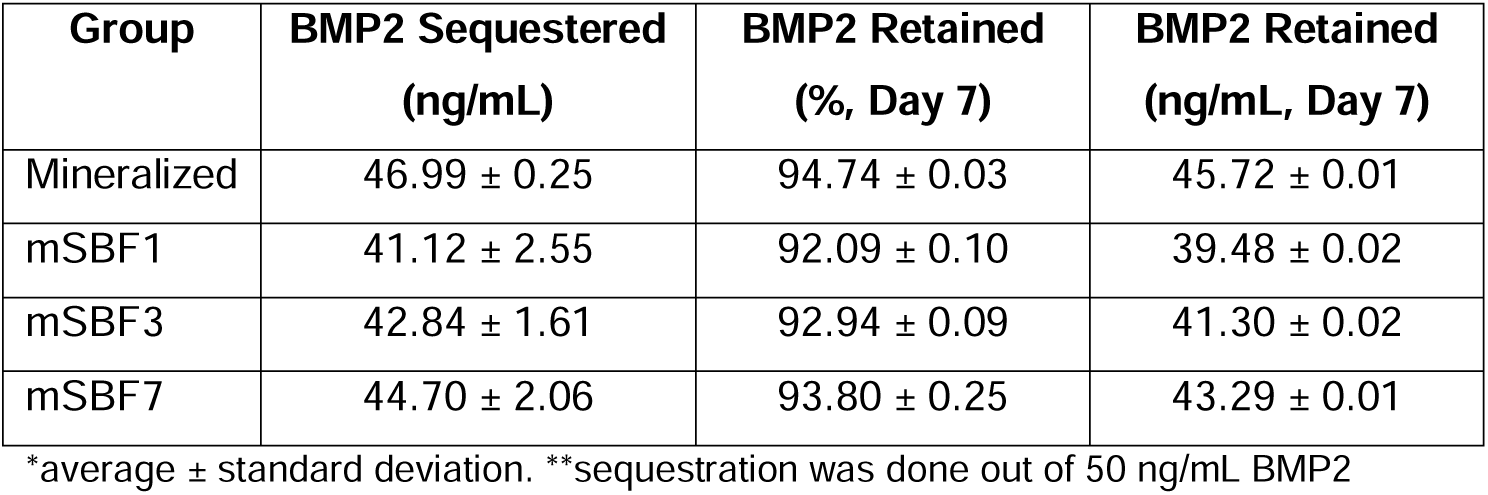
Sequestration and retention values for Figure 1 (mSBF treatments)

| Group | BMP2 Sequestered<br>(ng/mL) | BMP2 Retained<br>(%, Day 7) | BMP2 Retained<br>(ng/mL, Day 7) |
| --- | --- | --- | --- |
| Mineralized | 46.99 ± 0.25 | 94.74 ± 0.03 | 45.72 ± 0.01 |
| mSBF1 | 41.12 ± 2.55 | 92.09 ± 0.10 | 39.48 ± 0.02 |
| mSBF3 | 42.84 ± 1.61 | 92.94 ± 0.09 | 41.30 ± 0.02 |
| mSBF7 | 44.70 ± 2.06 | 93.80 ± 0.25 | 43.29 ± 0.01 |

\*average ± standard deviation. \*\*sequestration was done out of 50 ng/mL BMP2

Mechanical analysis of the different scaffolds showed the elastic modulus of scaffolds soaked in mSBF or PBS for 7 days is significantly lower than immediately after hydration (**Supplemental Figure 2**) with all groups softer than 5 kPa.

### 3.2. Sequential sequestration of individual growth factors increases retention in mineralized collagen scaffolds

We subsequently examined the use of mSBF treatments and sequential sequestrations to increase incorporation and retention of growth factors within our scaffolds compared to a single sequestration (one-time treatment). BMP2 and vascular endothelial growth factor (VEGF) can be sequentially sequestered into mineralized collagen scaffolds, and more VEGF was sequestered into mineralized collagen scaffolds than BMP2 for all treatment groups (**Figure 2B/D**). Notably, sequential sequestrations with repeated mBSF soaks resulted in significantly higher final concentrations of incorporated BMP2 than sequential sequestrations with repeated PBS soaks (**Figure 2B**). One-time treatments and the sequentially sequestered mSBF groups sequestered ∼72% and 66% of BMP2 out of solution, respectively, while sequentially sequestered PBS groups had significantly lower sequestration (∼63% BMP2 sequestered; **Figure 2B**). One-time treatments sequestered significantly more VEGF out of solution than both sequentially sequestered groups (mSBF and PBS; **Figure 2D**). One-time treatments sequestered ∼91% VEGF out of solution while both sequentially sequestered groups (mSBF and PBS) sequestered ∼82% VEGF out of solution (**Figure 2D**).

Next, we examined whether the mode of growth factor sequestration into the scaffold altered factor release. Notably, mineralized collagen scaffolds have higher retention of sequestered BMP2 when sequentially sequestered compared to a one-time treatment for the entire release period (**Figure 2C**). Sequentially sequestered PBS groups have higher BMP2 retention than sequentially sequestered mSBF groups by Day 7 (97.02% and 96.77%, respectively; **Figure 2C**). Although the sequentially sequestered groups have higher retention of sequestered BMP2, the final concentration of BMP2 in the scaffolds is highest in the one-time treatment group (54.77 ng/mL; **Table 2**). Mineralized collagen scaffolds with a one-time treatment and sequentially sequestered scaffolds with mSBF soaks have the highest retention of sequestered VEGF by Day 7 (**Figure 2E**), though all groups have greater than 98% of sequestered VEGF retained after 7 days (**Figure 2E**). Here, one-time treatments have the highest final concentration of VEGF (∼72 ng/mL) compared to the sequentially sequestered groups (∼65 ng/mL VEGF for mSBF and PBS treatments; **Table 2**). Summary statistics and full-scale retention data for this experiment: **Table 2, Supplemental Figure 3**.

**Table 2:** Sequestration and retention values for Figure 2 (individual growth factors)

| Group | BMP2 |  |  | VEGF |  |  |
| --- | --- | --- | --- | --- | --- | --- |
|  | Sequestered<br>(ng/mL) | Retained<br>(%, Day 7) | Retained<br>(ng/mL, Day 7) | Sequestered<br>(ng/mL) | Retained<br>(%, Day 7) | Retained<br>(ng/mL, Day 7) |
| One Trt | 57.90 ± 2.85 | 94.59 ± 0.03 | 54.77 ± 0.02 | 73.50 ± 0.44 | 98.79 ± 0.01 | 72.61 ± 0.01 |
| PBS | 51.06 ± 0.46 | 97.02 ± 0.02 | 49.54 ± 0.01 | 66.33 ± 0.16 | 98.42 ± 0.01 | 65.28 ± 0.01 |
| mSBF | 52.73 ± 0.26 | 96.77 ± 0.03 | 51.02 ± 0.01 | 66.34 ± 0.12 | 98.79 ± 0.1 | 65.53 ± 0.01 |

\*average ± standard deviation. \*\*sequestration was done out of 80 ng/mL BMP2 or 80 ng/mL VEGF

### 3.3. Sequential sequestration of multiple growth factors increases sequestration and retention in mineralized collagen scaffolds

We then explored whether the sequential sequestration approach provides an advantage for selectively incorporating multiple growth factors within the scaffold. Scaffolds could be exposed to a large dose of mixed factors (one-time treatments), or scaffolds could be sequentially exposed to one factor, followed by a PBS or mSBF treatment, then exposed to a second factor. Scaffolds that sequentially sequestered BMP2 then VEGF had higher concentrations of both growth factors compared to scaffolds that were soaked once in a solution containing BMP2 and VEGF (**Figure 3B/D**). One-time treatments sequestered ∼60% of both BMP2 and VEGF while sequentially sequestered groups sequestered ∼87% of BMP2 and greater than 90% of VEGF from solution. Data show it’s possible to quantify growth factor sequestration following each stage of exposure: (1) exposure to 40 ng/mL BMP2; (2) exposure to a second dose of 40 ng/mL BMP2; (3) exposure to 10 ng/mL VEGF (**Figure 3B/D**).

Sequentially sequestered groups (mSBF and PBS) also showed higher retention of both growth factors over 7 days when compared to one-time treatment scaffolds (**Figure 3C/E**). Sequentially sequestered scaffolds retained ∼98% and ∼95% of sequestered BMP2 and VEGF by Day 7, respectively. One-time treatments retained ∼94% and ∼88% BMP2 and VEGF by Day 7, respectively. Scaffolds that sequentially sequestered growth factors from solution had the highest final concentration of BMP2 and VEGF by Day 7 (∼68 ng/mL and ∼9 ng/mL, respectively; **Table 3**). Additionally, scaffolds that sequentially sequestered growth factors were less variable in their retention profiles (indicated by a narrower boxplot). Summary statistics and full-scale retention data for this experiment: **Table 3, Supplemental Figure 4**.

**Table 3:** Sequestration and retention values for Figure 3 (multiple growth factors)

| Group | BMP2 |  |  | VEGF |  |  |
| --- | --- | --- | --- | --- | --- | --- |
|  | Sequestered<br>(ng/mL) | Retained<br>(%, Day 7) | Retained<br>(ng/mL, Day 7) | Sequestered<br>(ng/mL) | Retained<br>(%, Day 7) | VEGF Retained<br>(ng/mL, Day 7) |
| One Trt | 50.64 ± 2.16 | 94.23 ± 1.62 | 47.72 ± 0.82 | 5.62 ± 0.43 | 88.95 ± 1.48 | 5.00 ± 0.08 |
| PBS | 71.91 ± 0.09 | 98.09 ± 0.08 | 70.53 ± 0.06 | 9.34 ± 0.07 | 94.25 ± 0.46 | 8.76 ± 0.04 |
| mSBF | 69.58 ± 0.06 | 98.04 ± 0.16 | 68.22 ± 0.11 | 9.24 ± 0.07 | 94.59 ± 0.18 | 8.75 ± 0.02 |

\*average ± standard deviation. \*\*sequestration was done out of 80 ng/mL BMP2 and 10 ng/mL VEGF

### 3.3. In vitro cell activity data suggest large growth factor doses are required to illicit cell response

We subsequently examined the ability for 10 ng/mL sequestered VEGF to induce shifts in human umbilical vein endothelial cell (HUVEC) activity using a transwell assay (i.e. examining the effect of *released* VEGF). We compared results for HUVECS in the scaffold with no exposure to VEGF (Blank), exposure to 10ng/mL VEGF supplemented in the media (Soluble), a single VEGF sequestration (One Trt), or sequential VEGF sequestration with mSBF treatments (mSBF). We did not observe significant differences in HUVEC metabolic activity between treatment groups in response to 10 ng/mL VEGF by Day 7 (**Supplemental Figure 5**). While we observed some short-term differences (Days 1 and 4) in gene expression for platelet derived factor (PDGF), hypoxia induced factor 1 alpha (HIF1A), angiopoietin-1 (ANG1), and angiopoietin-2 (ANG2), there was no significant effect of VEGF delivery method on long-term gene expression (Day 7) (**Supplemental Figure 6**).

## 4. Discussion

Bone healing is a complex process coordinated through growth factor signaling^21–25^, and tissue engineers use these factors to improve bone healing in bone-mimetic scaffolds^19,20^. Bone morphogenic protein 2 (BMP2)^20,42^, vascular endothelial growth factor (VEGF)^20^, and platelet derived growth factor (PDGF)^34^ are popular candidates for use in bone biomaterials. In this manuscript, we described the use of modified simulated body fluid (mSBF) and sequential sequestrations to incorporate BMP2 and VEGF into mineralized collagen scaffolds. This work seeks to adapt promising results using mSBF to deposit a mineral layer that can sequester growth factors onto surfaces^58–64^ to selectively incorporate growth factors used for bone repair applications into three-dimensional, porous biomaterials. Our primary goal is to improve growth factor incorporation and retention into mineralized collagen scaffolds via sequential sequestration strategies to enhance *in vivo* bone healing.

First, we hypothesized that extended exposure to mSBF prior to sequestration would increase growth factor incorporation and retention in mineralized collagen scaffolds. We found that 1- and 3-day mSBF treatments prior to sequestration *reduced* the amount of bone morphogenic protein 2 (BMP2) incorporated and retained in mineralized collagen scaffolds. This suggests the native mineral content of the mineralized collagen scaffolds is enough to sequester growth factors without additional treatments and that mSBF treatments prior to sequestration do not improve retention. Our mineralized collagen scaffolds are high in mineral content (40 weight percent^65^) and highly porous (>85%^66^), so it is likely that these two properties allowed for the high sequestration efficiency in our mineralized collagen scaffolds. We are able to control the mineral content^65^ and pore size^66^ of our materials, and future efforts will explore the effects of these characteristics on the sequestration and retention capabilities of mineralized collagen scaffolds. Scaffolds soaked in mSBF and PBS for 7 days were significantly softer than hydrated scaffolds, consistent with prior data showing that the osteogenic nature of the mineralized collagen scaffold is enhanced via release of mineral ions into the media^47^. While the porous nature of the scaffolds is essential for biological activity (i.e. cell penetration, diffusive biotransport), all scaffold variants show sub-optimal mechanical performance. While beyond the scope of this project, we have described methods to increase the mechanical stability of a cell activity optimized scaffolds via the inclusion of 3D printed polymeric meshes^49,69^. Based on these results, we did not use extended mSBF treatments prior to sequestration in subsequent experiments and instead focused on repeated mSBF treatments during sequestration.

Second, we hypothesized that sequential exposure to mSBF during growth factor sequestration would increase the incorporation and retention of growth factors in mineralized collagen scaffolds. Clements *et al.* has previously identified repeated mSBF treatments as a method to increase growth factor incorporation, maintain protein activity^60^, and prolong the release of growth factors^62^. One-time treatments and sequentially sequestered mSBF groups showed equal ability to incorporate BMP2 into the mineralized collagen scaffolds, but sequestered BMP2 had higher retention in the sequentially sequestered scaffolds (mSBF and PBS). This suggests that the mSBF and PBS soaks between sequestrations aid in the retention of BMP2 within mineralized collagen scaffolds. However, one-time treatments sequestered higher concentrations of VEGF out of solution than both sequentially sequestered groups, and sequestered VEGF had the highest retention in scaffolds with a one-time treatment and scaffolds sequentially sequestered with mSBF soaks. This prolonged VEGF release may be useful for *in vivo* implants, as suggested by previous work^45^.

VEGF (20-22 kDA; R&D systems catalog #293-VE) is approximately 37% larger than BMP2 (15-16 kDA; R&D systems catalog #355-BM), and this size difference might play a role in how these two factors are incorporated and maintained inside our mineralized collagen scaffolds. While we observed largely consistent results between mSBF and PBS soaks, we plan to test protein integrity and stability via circular dichroism^70,71^ or differential scanning calorimetry^72,73^. Additionally, we can confirm protein activity by evaluating the activation of receptors for the growth factor of interest in cells (e.g. vascular endothelial growth factor receptor, VEGFR^74,75^). Previous work utilizing mSBF treatments during sequestration suggests that the mineral coatings stabilize the growth factors and prolong their activity^62^. Such experiments may refine our understanding of whether mSBF soaks are necessary for extended protein activity in our scaffolds or if PBS soaks would be enough due to the already high mineral content in our materials^65,66^. Regardless, these results demonstrate that strategies to incorporate growth factors within the scaffold may require different methodologies to maximize incorporation and retention. As a result, it was essential to show the sequestration-based approach reported here could be adapted to sequentially incorporate multiple factors.

Third, we hypothesized that sequestering multiple growth factors via sequential sequestration would result in higher incorporation and retention of both factors. These results appear most promising because scaffolds that sequentially sequestered BMP2 then VEGF had significantly higher incorporation of both growth factors than scaffolds treated in a mixed solution of BMP2 and VEGF. Here, the one-time treatment group sequestered ∼60% of both BMP2 and VEGF from solution while sequentially sequestered groups sequestered ∼90% of BMP2 and VEGF from solution. Retention of both factors and final BMP2 and VEGF concentrations within the scaffolds was higher in the sequentially sequestered groups, meaning our mineralized collagen scaffolds sequester high concentrations of multiple factors within a single construct and have those factors retained for an extended period. We believe this sustained release would be particularly useful for cell migration and new vessel formation. Prolonged VEGF and platelet derived growth factor (PDGF) presence has been shown to promote migration, proliferation, and angiogenesis^45,76^.

Lastly, we evaluated cell activity in response to 10 ng/mL VEGF delivered through a transwell membrane (i.e. VEGF *released* from the scaffolds). These preliminary results suggest larger growth factors doses will be required to illicit extended cell responses (**Supplemental Figures 5, 6**). There are no significant effects of VEGF (soluble or sequestered) on human umbilical vein endothelial cell metabolic activity (**Supplemental Figure 5**) and gene expression (**Supplemental Figure 6**) compared to blank scaffolds (no VEGF) after 7 days. This is not surprising given the wide range of VEGF doses used *in vitro* and *in vivo* for tissue engineering purposes (25 ng/mL – 250 ug/mL)^45,77–80^, suggesting future efforts are required to define the correct dose for our mineralized collagen scaffolds. Immediate next steps are to determine the correct concentration of *released* growth factor needed to illicit a cellular response. However, we suspect that cells will be most influenced when cultured directly on the scaffolds due to the high concentration of growth factors retained within the material. Therefore, we will also culture cells within the scaffolds and analyze gene and protein expression to see how biologically active *retained* growth factors are. In both cases, released and retained growth factors, we are interested in dosing our mineralized collagen scaffolds with PDGF to induce cell migration and VEGF to induce vessel formation.

## 5. Conclusions

We have previously reported the use of covalent immobilization, photopatterning, and supramolecular interactions using cyclodextrins and glycosaminoglycans to incorporate growth factors into non-mineralized and mineralized collagen scaffolds. Here we explored sequential sequestration and the use of modified simulated body fluid to incorporate growth factors into mineralized collagen scaffolds and prolong growth factor release *in vitro*. We report the native mineralized collagen scaffolds can sequester growth factors (>90%) without any additional treatments, and this is a feature of these materials we have not previously explored. We show improved bone morphogenic protein 2 retention in mineralized collagen scaffolds using modified simulated body fluid treatments and sequential sequestrations. Notably, we demonstrate sequential sequestration can significantly increase incorporation and prolong retention for multi-factor cocktails (bone morphogenic protein 2, vascular endothelial growth factor). These methods provide an additional means to add growth factors into our materials and add complexity to aid in the healing of large bone injuries *in vivo*. Future work will investigate growth factor activity after sequestration and its influence on cellular activities critical to bone healing such as cell migration and vessel formation.

## Acknowledgements

This work was supported by the Office of the Assistant Secretary of Defense for Health Affairs Broad Agency Announcement for Extramural Medical Research through the Award No. W81XWH-16-1-0566. Opinions, interpretations, conclusions and recommendations are those of the authors and are not necessarily endorsed by the Department of Defense. Research reported in this publication was also supported by the National Institute of Dental and Craniofacial Research of the National Institutes of Health under Award Number R21 DE026582. The content is solely the responsibility of the authors and does not necessarily represent the official views of the NIH. We are grateful for the funding for this study provided by the NSF Graduate Research Fellowship DGE-1144245 (AST) and DGE-1144245 (MJD).

The authors would like to acknowledge the University of Illinois Roy J. Carver Biotechnology Center for assistance with real-time PCR. This research was carried out in part at the Imaging Technology Group within the Beckman Institute for Advanced Science and Technology at the University of Illinois at Urbana-Champaign. The authors would like to thank Scott Robinson and Cate Wallace for assistance with critical point drying and scanning electron microscopy. Additional support was provided by the Chemical and Biomolecular Engineering Department and the Carl R. Woese Institute for Genomic Biology (BACH) at the University of Illinois at Urbana-Champaign.

