## Supplemental Information for "Sequential sequestrations increase the incorporation and retention of multiple growth factors in mineralized collagen scaffolds"

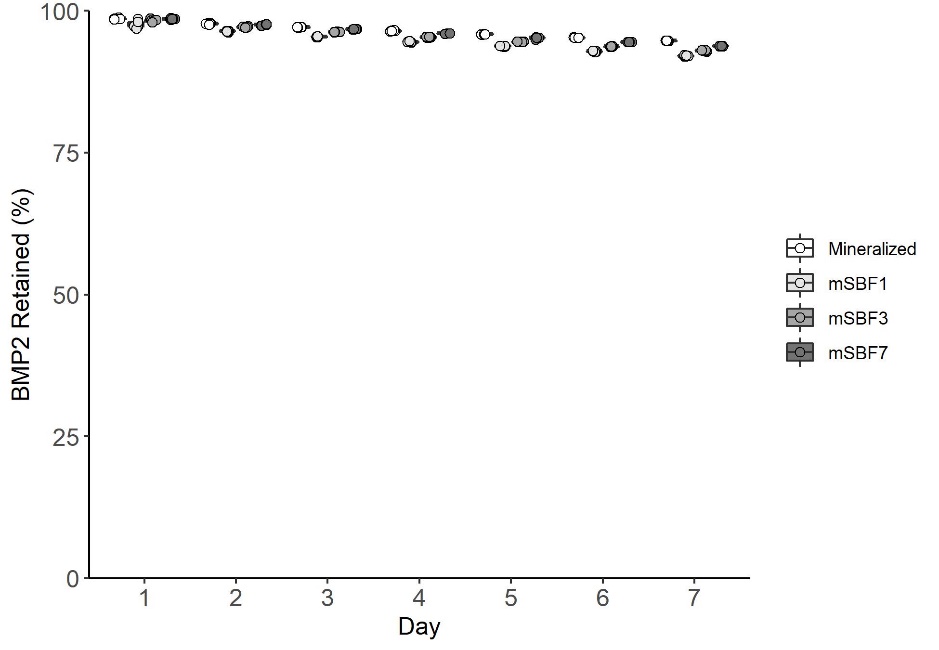

**Supplemental Figure 1: Full BMP2 retention plot for data presented in Figure 1C.** Retention of BMP2 in scaffolds with the y-axis starting at 0%.

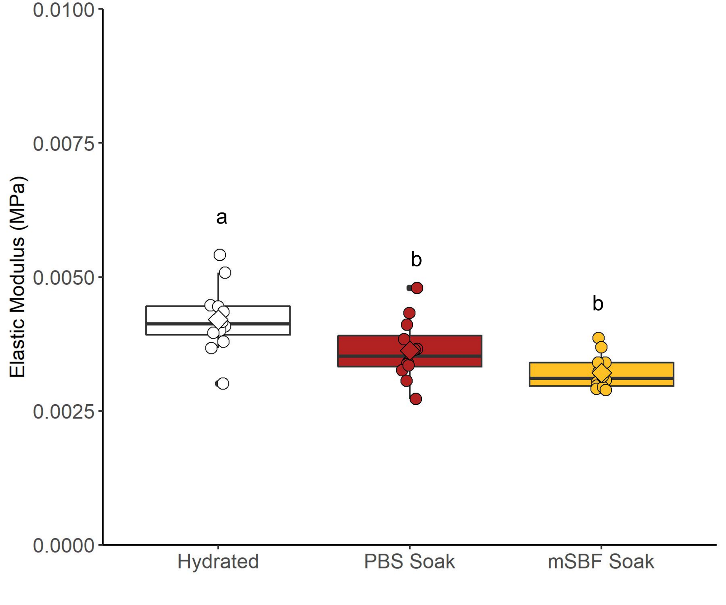

**Supplemental Figure 2: Compression testing of hydrated, mSBF soaked, and PBS soaked scaffolds.** (A) Elastic modulus of treated scaffolds. All scaffolds groups are softer than 5 kPa. Groups that share a letter are not significantly different (p<0.05).

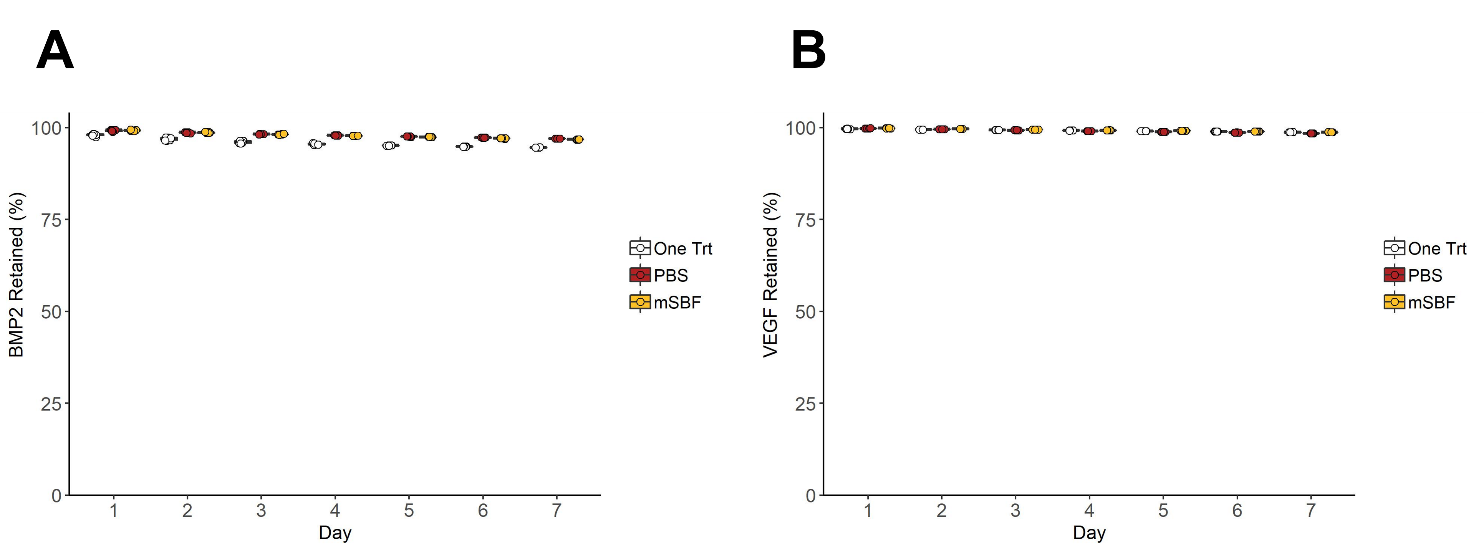

**Supplemental Figure 3: Full retention plots for data presented in Figure 2.** (A) Retention of BMP2 in scaffolds with the y-axis starting at 0%; corresponding to data presented in Figure 2C. (B) Retention of VEGF in scaffolds with the y-axis starting at 0%; corresponding to data presented in Figure 2E.

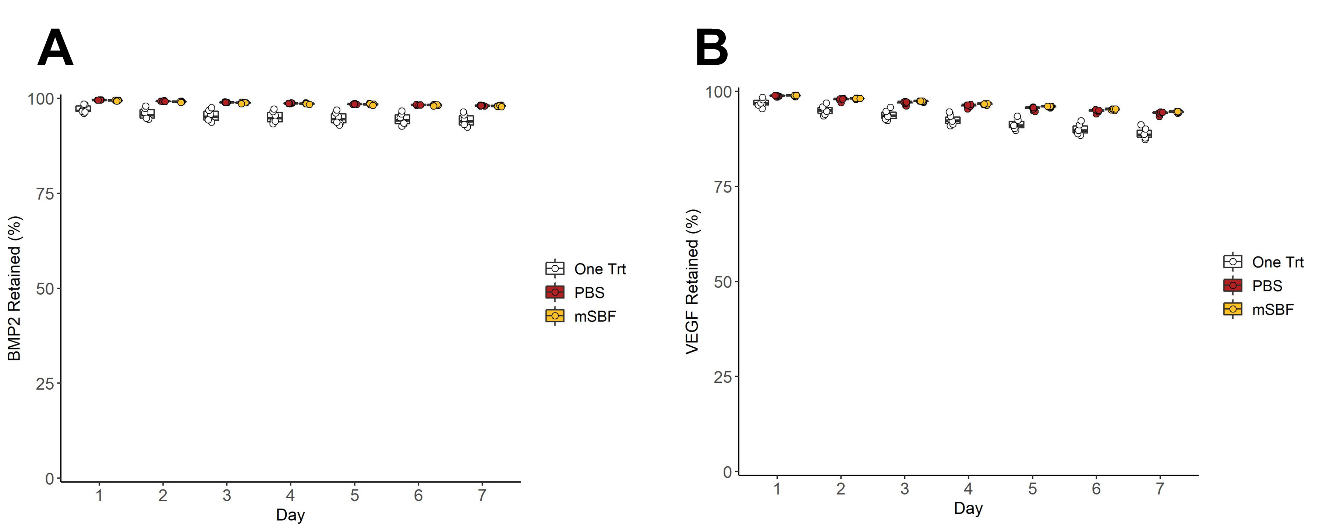

**Supplemental Figure 4: Full retention plots for data presented in Figure 3.** (A) Retention of BMP2 in scaffolds with the y-axis starting at 0%; corresponding to data presented in Figure 3C. (B) Retention of VEGF in scaffolds with the y-axis starting at 0%; corresponding to data presented in Figure 3E.

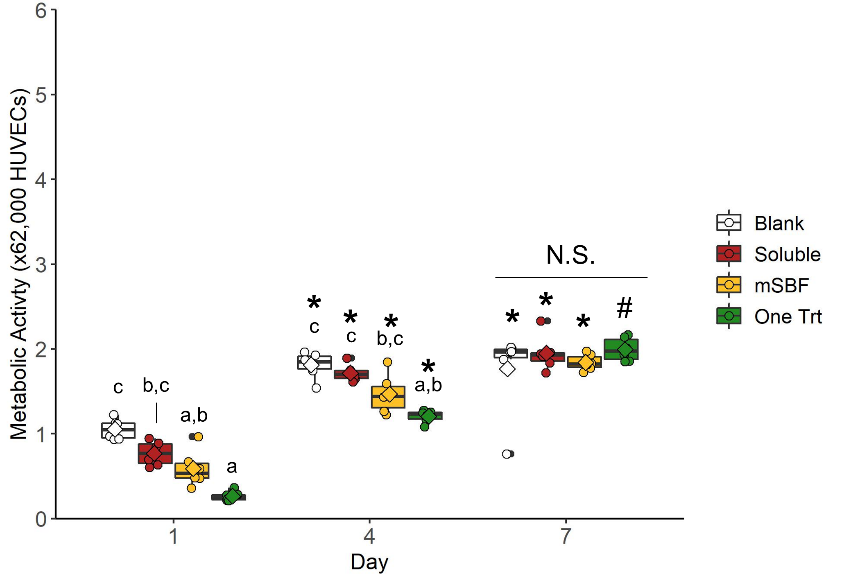

**Supplemental Figure 5: Cell metabolic activity in response to vascular endothelial growth factor released from scaffolds.** There is no significant difference in human umbilical vein endothelial cell metabolic activity between groups by Day 7. Groups that share a letter within a day are not significantly different (p<0.05). N.S. indicates no significant difference between indicated groups (p<0.05). * indicates significance compared to the same group at Day 1 (p<0.05). # indicates significance compared to the same group at Day 1 and Day 4 (p<0.05).

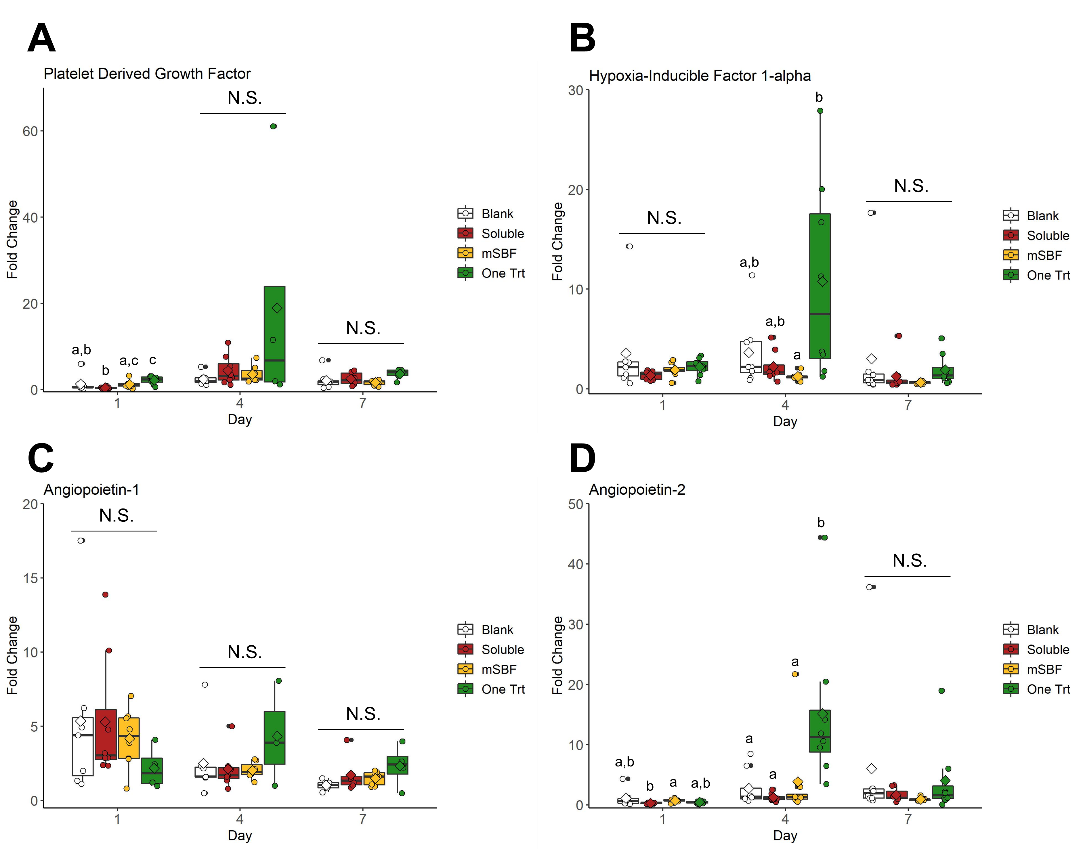

**Supplemental Figure 6: Gene expression in response to vascular endothelial growth factor released from scaffolds.** (A) Platelet derived growth factor fold change. (B) Hypoxia induced factor 1 alpha fold change. (C) Angiopoietin-1 fold change. (D) Angiopoietin-2 fold change. Overall, there are no significant differences in human umbilical vein endothelial cell gene expression between treatments groups by Day 7. Groups that share a letter within a day are not significantly different (p<0.05). N.S. indicates no significant difference between indicated groups (p<0.05).

**Supplemental Table 1.** PCR primers and assay IDs.

| **Transcript** | **Supplier** | **Assay ID** |
| --- | --- | --- |
| *18S* | ThermoFisher (Taqman) | Hs99999901_s1 |
| *PPIA* | ThermoFisher (Taqman) | Hs04194521_s1 |
| *PDGF* | ThermoFisher (Taqman) | Hs00966522_m1 |
| *HIF1a* | ThermoFisher (Taqman) | Hs00153153_m1 |
| *ANG1* | ThermoFisher (Taqman) | Hs00919202_m1 |
| *ANG2* | ThermoFisher (Taqman) | Hs00169867_m1 |

**Supplemental Table 2.** Summary statistics for PDGF and HIF1a PCR data

|  |  | **Platelet Derived Growth Factor** | | **Hypoxia Inducible Factor 1α** | |
| --- | --- | --- | --- | --- | --- |
| **Day** | **Group** | **Fold Change** | **Sample Size** | **Fold Change** | **Sample Size** |
| 1 | Blank | 1.310 ± 2.086 | 7 | 3.565 ± 4.796 | 7 |
| 1 | mSBF | 1.246 ± 0.902 | 8 | 1.902 ± 0.696 | 8 |
| 1 | One Trt | 2.191 ± 0.921 | 7 | 2.208 ± 0.845 | 8 |
| 1 | Soluble | 0.440 ± 0.199 | 8 | 1.363 ± 0.417 | 8 |
| 4 | Blank | 2.461 ±1.381 | 7 | 3.645 ± 3.468 | 8 |
| 4 | mSBF | 3.592 ± 2.005 | 7 | 1.248 ± 0.379 | 8 |
| 4 | One Trt | 18.95 ± 28.48 | 4 | 10.76 ± 9.932 | 8 |
| 4 | Soluble | 4.430 ± 3.353 | 8 | 2.226 ± 1.506 | 8 |
| 7 | Blank | 2.123 ± 2.003 | 8 | 3.014 ± 5.937 | 8 |
| 7 | mSBF | 1.624 ± 0.654 | 6 | 0.647 ± 0.126 | 8 |
| 7 | One Trt | 3.619 ± 1.184 | 5 | 1.916 ± 1.561 | 8 |
| 7 | Soluble | 2.514 ± 1.543 | 6 | 1.249 ± 1.669 | 8 |

**Supplemental Table 3.** Summary statistics for ANG1 and ANG2 PCR data

|  |  | **Angiopoietin 1** | | **Angiopoietin 2** | |
| --- | --- | --- | --- | --- | --- |
| **Day** | **Group** | **Fold Change** | **Sample Size** | **Fold Change** | **Sample Size** |
| 1 | Blank | 5.360 ± 5.702 | 7 | 1.104 ± 1.462 | 7 |
| 1 | mSBF | 4.176 ± 1.991 | 8 | 0.737± 0.290 | 8 |
| 1 | One Trt | 2.182 ± 1.427 | 4 | 0.455 ± 0.186 | 8 |
| 1 | Soluble | 5.309 ± 4.315 | 8 | 0.302 ± 0.086 | 8 |
| 4 | Blank | 2.508 ± 2.410 | 7 | 2.742 ± 2.994 | 8 |
| 4 | mSBF | 2.030 ± 0.550 | 7 | 3.865 ± 7.248 | 8 |
| 4 | One Trt | 4.324 ± 3.557 | 3 | 15.12 ± 12.86 | 8 |
| 4 | Soluble | 2.126 ± 1.365 | 7 | 1.251 ± 0.656 | 8 |
| 7 | Blank | 1.035 ± 0.335 | 6 | 6.024 ± 12.18 | 8 |
| 7 | mSBF | 1.495 ± 0.447 | 8 | 0.936 ± 0.286 | 8 |
| 7 | One Trt | 2.335 ± 1.438 | 4 | 4.058 ± 6.275 | 8 |
| 7 | Soluble | 1.718 ± 1.195 | 6 | 1.666 ± 1.115 | 7 |
